## Supplemental data, tables, and figures for "Sliding of HIV-1 reverse transcriptase over DNA creates a transient P pocket – Targeting P-pocket by fragment screening"

**Contents**

Supplementary Movie legends 3

Supplementary Table 1. X-ray crystallography data collection and refinement 4

Supplementary Fig. 1. Crystal symmetry interaction creates P-pocket 5

Supplementary Fig. 2. A schematic overview of the experimental setups used in XChem facility to perform fragment screening by X-ray crystallography 6

Supplementary Table 2. Virtually designed fragments 7

Supplementary Table 3. Docking results and drug-like properties of hits and selected fragments 8

Supplementary Fig. 3. Docking mode and detailed interactions of designed fragments F01-F05 with HIV-1 RT/dsDNA 9

Supplementary Fig. 4. Interactions of (a) **F47** and (b) **F81** with HIV-1 RT/dsDNA 10

Supplementary Fig. 5. Size-exclusion chromatography of HIV-1 RT/37-mer hairpin-DNA aptamer complex 11

Supplementary Table 4. Single particle cryo-EM data and structure analysis statistics 12

Supplementary Fig. 6. Impacts of crystallography and cryo-EM experimental conditions on the complexes 13

Supplementary Fig. 7. Cryo-EM data processing 14

Fragment design and docking study 15

Synthesis of fragments (Schemes 1-3; Supplementary Figure 8) 17

NMR spectra and HRMS of fragments **F01 – F05** 20

References 28

### 1. Supplementary Movies

**Movie 1. Sliding of RT over a dsDNA substrate**. Sliding of RT over a dsDNA substrate. The structures of N, P, and P-1 complexes of I63C RT/DNA were morphed to show sliding of RT over a dsDNA. The movie shows the transition starting from the N complex (light gray) to P-1 complex (salmon) via P complex; in the movie, the position of RT is fixed, and the DNA is sliding. The fingers were closed in the N and P complexes as the N-site (dNTP-binding pocket) is occupied in those structures.

**Movie 2. A zoomed view of sliding at the polymerase active site.** The color code and sliding are as in Movie 1.

**Movie 3. Rearrangement of P-pocket to accommodate the fragment 166.** Morphing between the fragment 166-bound RT/DNA (salmon) and apo RT/DNA (light gray) structures shows the rearrangement of the P-pocket to bind the fragment.

**Supplementary Table 1. X-ray crystallography data collection and refinement statistics.**

| **Data collection** | **Apo RT/DNA** | **Fragment 048** | **Fragment 166** |
|---|---|---|---|
| Synchrotron Beamline | Diamond I04 | Diamond I04-1 | Diamond I04-1 |
| Wavelength (Å) | 0.91587 | 0.91589 | 0.91589 |
| Space group | *C*2 | *C*2 | *C*2 |
| Molecule/a.s.u. | 2 | 2 | 2 |
| Cell dimensions | | | |
| *a*, *b*, *c* (Å) | 310.63, 62.07, 168.23 | 309.57, 61.90, 168.78 | 310.74, 62.06, 169.29 |
| $\alpha, \beta, \gamma$ (°) | 90, 104.51, 90 | 90, 104.55, 90 | 90, 104.93, 90 |
| Resolution, Å | 81.44 – 2.85 | 98.75 – 3.30 | 150.13 – 3.37 |
| (highest resolution shell) | (2.91 – 2.85)* | (3.42 – 3.30) | (3.43 – 3.37) |
| Unique reflections | 73287 (4471) | 44516 (4374) | 44790 (2220) |
| $R_{merge}$ | 0.234 (1.413) | 0.323 (0.953) | 0.329 (2.88) |
| I / $\sigma$(I) | 5.9 (1.5) | 3.3 (1.4) | 3.4 (0.5) |
| $CC_{1/2}$ | 0.991 (0.243) | 0.973 (0.251) | 0.987 (0.338) |
| Completeness (%) | 99.8 (100) | 94.6 (95.3) | 99.9 (100) |
| Redundancy | 6.2 (6.2) | 3.4 (3.5) | 5.7 (5.9) |
| **Refinement** | | | |
| Resolution (Å) | 2.85 | 3.30 | 3.37 |
| $R_{work}$ / $R_{free}$ | 0.21/0.24 | 0.26/0.29 | 0.22/0.26 |
| No. atoms | 17865 | 17578 | 17725 |
| Macromolecules | 17669 | 17530 | 17613 |
| Ligand[#] | - | 26 | 29 |
| Water | 157 | - | 19 |
| *B*-factors (Å$^2$) | | | |
| Macromolecules | 63.86 | 63.27 | 98.62 |
| Ligand | - | 77.39 | 121.49 |
| Water | 44.73 | - | 73.50 |
| R.m.s. deviations | | | |
| Bond lengths (Å) | 0.003 | 0.005 | 0.005 |
| Bond angles (°) | 0.550 | 0.905 | 0.862 |
| Ramachandran plot | | | |
| Favoured/allowed/ outlier (%) | 96.97/2.87/0.16 | 96.56/3.18/0.26 | 96.87/2.97/0.16 |
| Rotamer | | | |
| Favoured/poor (%) | 80.99/0.75 | 91.88/0.29 | 91.49/0.17 |
| MolProbity scores | | | |
| Protein geometry | 1.55 (100$^{th}$) | 1.90 (100$^{th}$) | 1.84 (100$^{th}$) |
| Clash score all atoms | 6.86 (100$^{th}$) | 16.44 (97$^{th}$) | 13.96 (97$^{th}$) |
| PDB code | 7OZ2 | 7OXQ | 7OZ5 |

*values in parentheses are for highest-resolution shell.

[#]bound fragment.

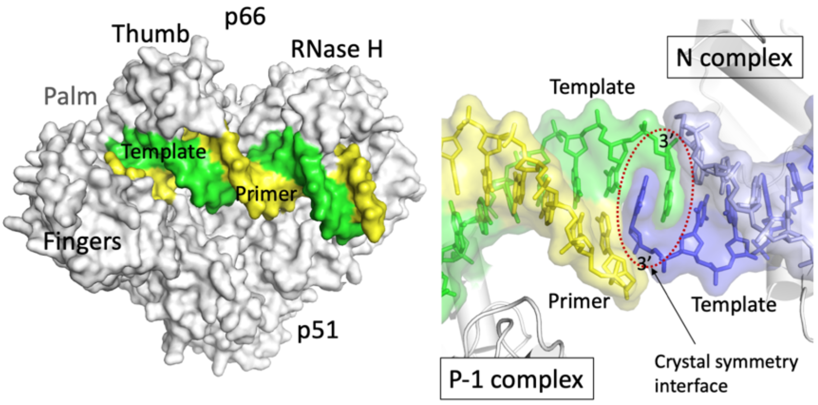


**Supplementary Fig. 1. Crystal symmetry interaction creates P-pocket. a** Space-filling model of RT (gray) with dsDNA (yellow primer, green template). **b** Crystal symmetry interaction between the DNA duplexes of P-1 complex (left; green and yellow) and N complex (right; dark and light blue) stabilizes the P-1 complex with a transient P-pocket in crystal.

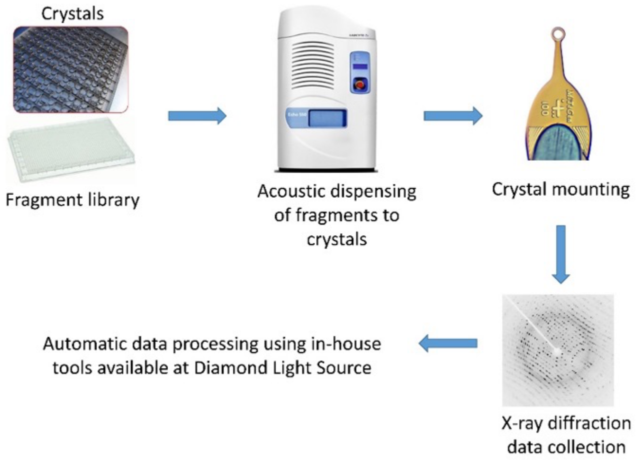


**Supplementary Fig. 2. A schematic overview of the experimental setups used at XChem facility for fragment screening by X-ray crystallography.** Fragment libraries were stored in 96 well plates; one fragment solution per well. An acoustic dispenser was used to dispense a small volume of a given fragment directly to a drop on a 96-well crystallization plate containing crystals in each well. After soaking, one crystal per fragment were mounted on cryogenic loops[4] flash frozen in liquid $N_2$ and loaded to the I04-1 beamline for automatic data collection.

**Supplementary Table 2**. **Virtually designed fragments.**

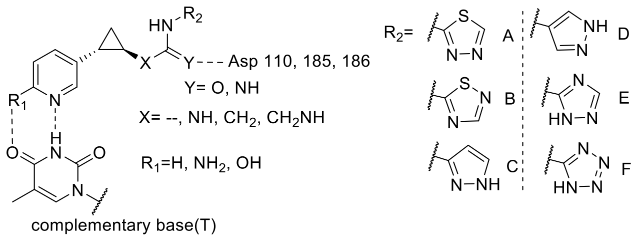

complementary base(T)

| Frag. | $R_1$ | $R_2$ | X | Y | Frag. | $R_1$ | $R_2$ | X | Y | Frag. | $R_1$ | $R_2$ | X | Y |
|---|---|---|---|---|---|---|---|---|---|---|---|---|---|---|
| F01 | H | A | -- | O | F29 | $NH_2$ | E | -- | NH | F57 | H | C | $CH_2$ | NH |
| F02 | H | B | -- | O | F30 | $NH_2$ | F | -- | NH | F58 | H | D | $CH_2$ | NH |
| F03 | H | C | -- | O | F31 | OH | A | -- | NH | F59 | H | E | $CH_2$ | NH |
| F04 | H | D | -- | O | F32 | OH | B | -- | NH | F60 | H | F | $CH_2$ | NH |
| F05 | H | E | -- | O | F33 | OH | C | -- | NH | F61 | $NH_2$ | A | $CH_2$ | NH |
| F06 | H | F | -- | O | F34 | OH | D | -- | NH | F62 | $NH_2$ | B | $CH_2$ | NH |
| F07 | $NH_2$ | A | -- | O | F35 | OH | E | -- | NH | F63 | $NH_2$ | C | $CH_2$ | NH |
| F08 | $NH_2$ | B | -- | O | F36 | OH | F | -- | NH | F64 | $NH_2$ | D | $CH_2$ | NH |
| F09 | $NH_2$ | C | -- | O | F37 | H | A | NH | NH | F65 | $NH_2$ | E | $CH_2$ | NH |
| F10 | $NH_2$ | D | -- | O | F38 | H | B | NH | NH | F66 | $NH_2$ | F | $CH_2$ | NH |
| F11 | $NH_2$ | E | -- | O | F39 | H | C | NH | NH | F67 | OH | A | $CH_2$ | NH |
| F12 | $NH_2$ | F | -- | O | F40 | H | D | NH | NH | F68 | OH | B | $CH_2$ | NH |
| F13 | OH | A | -- | O | F41 | H | E | NH | NH | F69 | OH | C | $CH_2$ | NH |
| F14 | OH | B | -- | O | F42 | H | F | NH | NH | F70 | OH | D | $CH_2$ | NH |
| F15 | OH | C | -- | O | F43 | $NH_2$ | A | NH | NH | F71 | OH | E | $CH_2$ | NH |
| F16 | OH | D | -- | O | F44 | $NH_2$ | B | NH | NH | F72 | OH | F | $CH_2$ | NH |
| F17 | OH | E | -- | O | F45 | $NH_2$ | C | NH | NH | F73 | H | A | $CH_2NH$ | NH |
| F18 | OH | F | -- | O | F46 | $NH_2$ | D | NH | NH | F74 | H | B | $CH_2NH$ | NH |
| F19 | H | A | -- | NH | F47 | $NH_2$ | E | NH | NH | F75 | H | C | $CH_2NH$ | NH |
| F20 | H | B | -- | NH | F48 | $NH_2$ | F | NH | NH | F76 | H | D | $CH_2NH$ | NH |
| F21 | H | C | -- | NH | F49 | OH | A | NH | NH | F77 | H | E | $CH_2NH$ | NH |
| F22 | H | D | -- | NH | F50 | OH | B | NH | NH | F78 | H | F | $CH_2NH$ | NH |
| F23 | H | E | -- | NH | F51 | OH | C | NH | NH | F79 | $NH_2$ | A | $CH_2NH$ | NH |
| F24 | H | F | -- | NH | F52 | OH | D | NH | NH | F80 | $NH_2$ | B | $CH_2NH$ | NH |
| F25 | $NH_2$ | A | -- | NH | F53 | OH | E | NH | NH | F81 | $NH_2$ | C | $CH_2NH$ | NH |
| F26 | $NH_2$ | B | -- | NH | F54 | OH | F | NH | NH | F82 | $NH_2$ | D | $CH_2NH$ | NH |
| F27 | $NH_2$ | C | -- | NH | F55 | H | A | $CH_2$ | NH | F83 | $NH_2$ | E | $CH_2NH$ | NH |
| F28 | $NH_2$ | D | -- | NH | F56 | H | B | $CH_2$ | NH | F84 | $NH_2$ | F | $CH_2NH$ | NH |

**Supplementary Table 3. Docking results and drug-like properties of hits and selected fragments.**

| Fragment No. | Chemical structure | Affinity (kcal/mol) | Number of H bond interactions[a] | M.W. | cLogP[b] | Interacting surface area (Å$^2$) of ligand[c] |
|---|---|---|---|---|---|---|
| 048 | 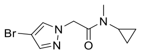 | -5.1 | 1 | 258.12 | 1.07 | 333.3 |
| 166 | 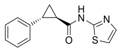 | -5.9 | 2 | 244.31 | 2.68 | 339.5 |
| F01 | 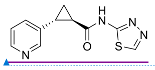 | -6.7 | 0 | 246.29 | 0.86 | 347.8 |
| F02 | 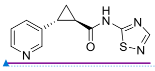 | -6.2 | 1 | 246.29 | 0.87 | 332.8 |
| F03 | 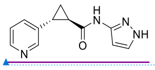 | -6.7 | 2 | 228.26 | 0.35 | 348.4 |
| F04 | 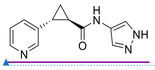 | -6.7 | 2 | 228.26 | 0.63 | 367.7 |
| F05 | 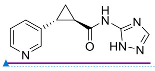 | -6.4 | 2 | 229.24 | 0.31 | 377.2 |
| F47 | 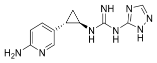 | -6.6 | 3 | 258.29 | -0.12 | 378.9 |
| F81 | 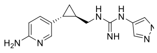 | -7.6 | 4 | 271.33 | 0.78 | 438.8 |

[a] Calculated from LigPlot.

[b] Calculated from http://www.vcclab.org/lab/alogps/.

[c] Generated from https://www.ebi.ac.uk/pdbe/pisa/

Field Code Changed
Field Code Changed
Field Code Changed
Field Code Changed
Field Code Changed

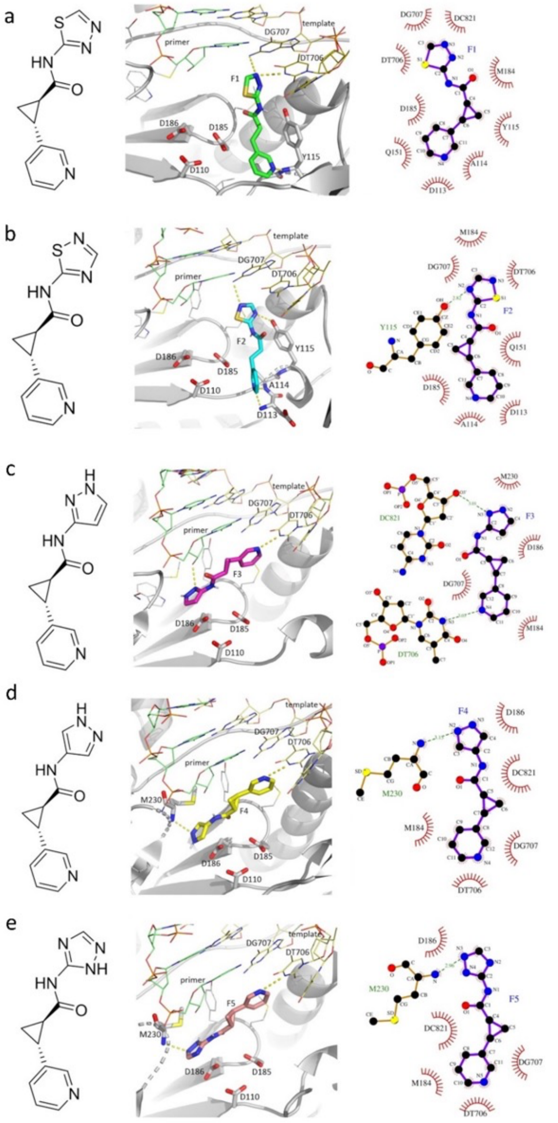


**Supplementary Fig. 3. Docking mode and detailed interactions of designed fragments F01-F05 with HIV-1 RT/dsDNA** (panel **a-e**). Conformation having best free energy of binding is shown in each case. Hydrogen bond (H-bond) interactions are shown with yellow dashed lines; binding poses were prepared in PyMol,[1] and detailed interaction diagrams were obtained with LigPlot+ v.2.2.[2]

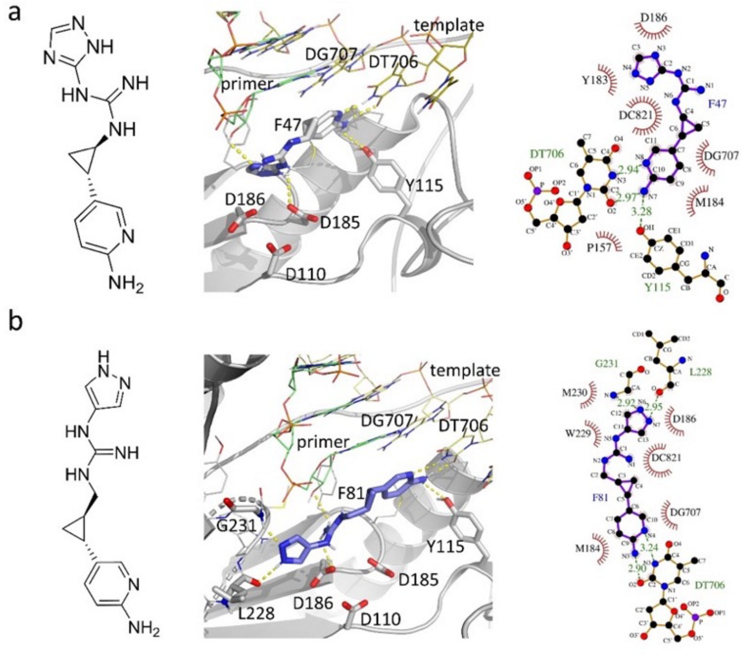


**Supplementary Fig. 4. Interactions of (a) F47 and (b) F81 with HIV-1 RT/dsDNA.** Conformation having best free energy of binding is shown in each case. H-bond interactions are depicted in yellow dashed lines; docking poses were generated in PyMol,[1] and detailed interaction diagrams were obtained with LigPlot+ v.2.2.[2]

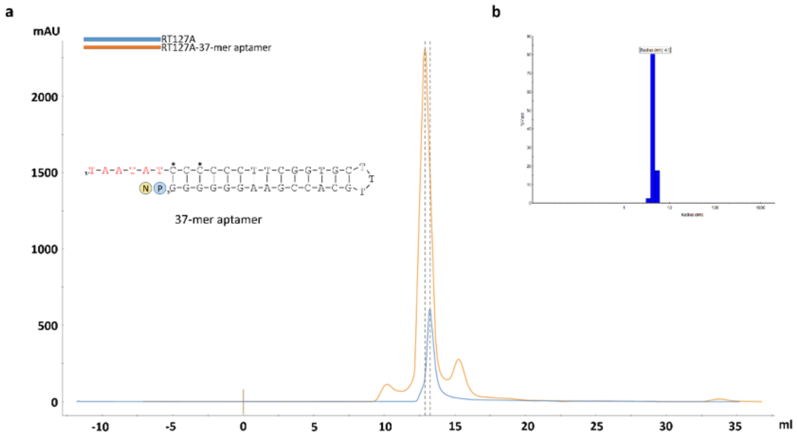


**Supplementary Fig. 5. Size-exclusion chromatography of HIV-1 RT/37-mer hairpin-DNA aptamer complex. a** sequence of the 37-mer hairpin-DNA aptamer. Template overhang is in red, P-site and N-site are shown under blue and yellow circles respectively. Eluted peak of the RT-aptamer complex (orange) is compared with that of the RT only peak (blue) to highlight the shift in the profile; mid points of the peaks are labelled with vertical dashed lines. **b** dynamic light scattering (DLS) profile of the RT-aptamer complex showing particle size and sample homogeneity.

**Supplementary Table 4. Single particle cryo-EM data and structure analysis statistics.**

| Structure | RT/Aptamer DNA/**166** | RT/Aptamer/**F04** |
|---|---|---|
| PDB ID/EMBD ID | 7OZW/EMD-13139 | 7P15/EMD-13156 |
| Data collection | | |
| Grid type | Quantifoil R1.2/1.3 | Quantifoil R1.2/1.3 |
| Number of grids | 1 | 1 |
| Microscope/detector | Glacios/Falcon 3 | Glacios/Falcon 3 |
| Voltage (kV) | 200 | 200 |
| Magnification | 150,000 x | 150,000 x |
| Recording mode | Counting | Counting |
| Dose ($e^-$/Å$^2$/frame) | 1.25 | 1.25 |
| Total dose (e/Å$^2$) | 50 | 50 |
| Number of frames/movies | 40 | 40 |
| Total exposure time (sec) | 55 | 55 |
| Pixel size (Å) | 0.97 | 0.97 |
| Defocus range (Å) | -8000 to - 18000 | -8000 to - 18000 |
| Data processing | | |
| Number of micrographs used | 660 | 767 |
| Number of particles picked | 856,171 | 733,223 |
| Particles used for final map | 146,670 | 157,094 |
| Fourier Completeness | 0.911 | 0.89 |
| Map resolution (FSC 0.143; Å) | 3.38 | 3.58 |
| Map sharpening B factor (Å$^2$) | 140.2 | 148.9 |
| Model fitting | | |
| Experimental map/model correlation | 0.70 | 0.72 |
| Experimental map/ligand correlation | 0.43 | 0.53 |
| Total number of atoms | 8,407 | 8,639 |
| Number of residues/Average B factor (Å$^2$) | | |
| Protein | 961/35.08 | 969/55.60 |
| Nucleic acid | 34/82.97 | 34/94.53 |
| Ligand | 1/65.50 | 1/63.73 |
| Clash score | 7.07 | 8.4 |
| Ramachandran plot; favored/outlier (%) | 96.34/0.0 | 96.05/0.0 |
| Rotamer outlier (%) | 0.0 | 0.12 |
| RMSD bond length (Å)/bond angle (°) | 0.004/0.74 | 0.004/0.62 |
| MolProbity score | 1.63 | 1.79 |

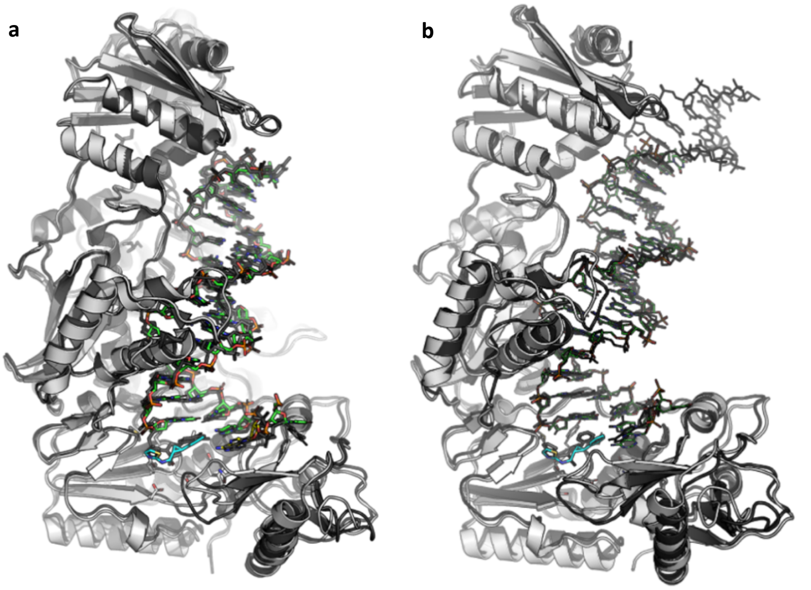


**Supplementary Fig. 6. Impacts of crystallography and cryo-EM experimental conditions on the complexes. a.** Superposition of cryo-EM structure of RT/37-aptamer DNA/166 complex (light gray RT, green DNA, and cyan **166**) and crystal structure of RT/38-aptamer DNA (PDB ID. 5D3G, dark gray) revealed that the structural features including the track of DNA aptamer, subdomain arrangements, and DNA-protein interactions are conserved between two structures. The fragment **166** is bound to P-pocket in P-1 complex cryo-EM structure, and the pocket is occupied by the 3'-end nucleotide of aptamer DNA in the crystal structure. A total of 866 Cα atoms superimposed with rmsd of 0.95 Å. **b.** The cryo-EM structure of RT/37-aptamer DNA/**166** complex (light gray RT, green DNA, and cyan **166**) and crystal structure of cross-linked RT/DNA/**166** complex (dark gray) superimpose well with rmsd of 1.14 Å for 931 aligned C$\alpha$ atoms.

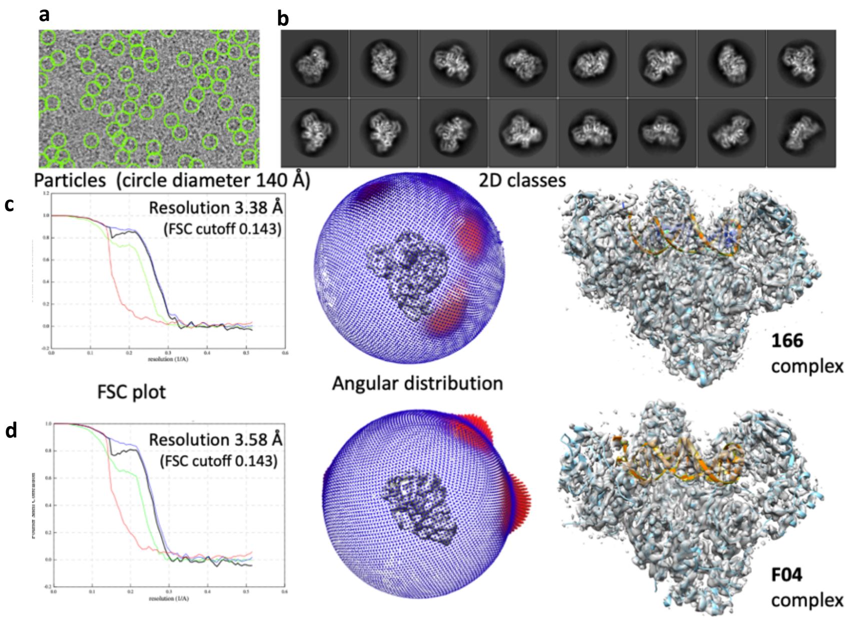


**Supplementary Fig. 7. Cryo-EM data processing. a.** Picked particles on a typical micrograph of RT/aptamer-DNA/166 complex. **b.** Selected 2D classes. FSC resolution, angular distribution of particles in the final set, and B-sharpened map covering the model for RT/aptamer-DNA/**166** complex structure (row **c**) and RT/aptamer-DNA/**F04** complex structure (row **d**).

**Fragment design and docking study**

Creation of a well-formed P-pocket and the binding of the fragments **048** and **166** to the pocket intuited the need for further analysis of the pocket characteristics. We attempted virtually designing a set of compounds with the chemical backbone of **166**, such that the analogs can form H-bond with template thymine base. Therefore, the phenyl ring was replaced by a pyridine moiety. In addition, an amino or hydroxyl group was introduced at the 5'-position of the pyridine ring so that two H-bonds could be potentially formed like base-pairing. At the same time, we maintained the amide bond of **166**, and made variations of the thiazole moiety with different five-membered heteroaromatics. Eighteen (**F01-F18**) analogs of **166** were modeled by accommodating the above discussed variations (Supplementary **Table 1**).

In order to investigate the favorability of binding of the designed fragments to P-pocket, we conducted molecular docking study; initially rigid docking followed by flexible docking using Autodock Vina (ver. 1.1.1).[3] For flexible docking, we defined the catalytic residues D110, D185 and D186 of HIV-1 RT as flexible residues. Analysis of docking results revealed that ten fragments (**F03-F08, F10-F13**) have the potential for forming H-bonds with template thymine overhang. In addition, the newly introduced heteroaromatics can engage in additional H-bond formation with surrounding residues and DNA primer. Specifically, fragments **F03, F8, F10-11** formed one H-bond with 3'-hydroxyl group of the primer end nucleotide DC821, while **F04-F06** formed one H-bond with backbone NH of primer grip residue M230. Moreover, **F07, F12-13** formed H-bonds with both M230 and the primer 3'-end nucleotide. However, all above-mentioned fragments did not acquire H-bond interactions with catalytic residues D110, D185 and/or D186. Based on these observations and taking into account the synthesis feasibility, we proceeded to synthesize fragments **F01-F05** whose docking outcomes are shown (Supplementary Figure 1, Supplementary Table 2).

The guanidinium group as present in the side-chain of arginine interacts with carboxylate groups via salt bridges, which can be found in many crystal structures of enzyme complexes with oxoanionic substrates and simple guanidinium salts.[4] To explore the possibility of our compounds forming such interactions with the catalytic residues, we decided to substitute the amido linkage for an amidine or guanidine linker and make small variation on the linker length to increase flexibility, resulting in the design of the fragments **F19-F84** (Supplementary Table 1).

Molecular docking study demonstrated that fragments **F19-F24** with an amidine linker can form H-bonds with the template thymine and with the residue D185, whereas **F26-F30** form H-bonds with thymine only. Fragments (**F43, F45-46** and **F52**) with a guanine linker can only form H-bonds with template thymine, whereas **F47** and **F48** can also interact with D185. Interestingly, although fragments (F62-F64) with longer guanine linker only form H-bonds with thymine, their counterparts (F60, F66, F79-F84) can also form one additional interaction with the residue D186. For these fragments, the newly introduced heteroaromatics (Supplementary Table 1) can also form H-bond with the primer grip/primer 3'-nucleotide. Both **F81** and **F82** can form one H-bond with 3'-hydroxyl end of the primer strand and backbone of residue L228, respectively, and the former can form an additional H-bond with G231 while maintaining interaction with the template thymine.

Considering the binding affinity, stability and structural variation of designed fragments, and ease of synthesis we selected two fragments (**F47** and **F81**) for synthesis. The detailed docking results and drug-like properties of hits (**048** and **166**) and selected fragments are shown in Supplementary Table 2. Docking score of selected fragments were improved when compared with that of two fragment screen hits. In addition, the docked modes of **F47** and **F81** suggested more interacting surface areas than that of two hits (Supplementary Fig. 2). Furthermore, selected fragments satisfy Lipinski's Rule of Five.[5]

**Synthesis of fragments**

In view of their convenient synthesis, we firstly prepared **F01-05** for structural study to examine if the pyridyl ring can form H-bond with template thymine. The preparation of these compounds is depicted in **Scheme 1**. The *trans*-cyclopropane-containing compounds were all synthesized as racemic mixtures.

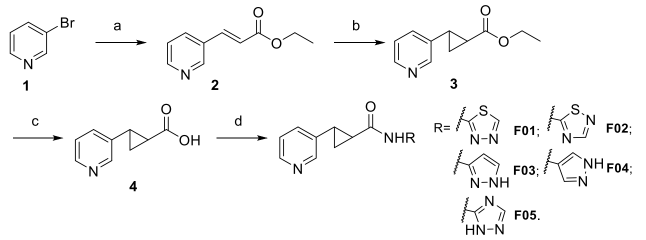


**Scheme 1.** *Reagents and conditions*: (a) Ethyl acrylate, $Pd(OAc)_2$, $K_2CO_3$, $PPh_3$, DMF, 100 °C, 20 h, 75 %; (b) $Me_3SOI$, NaH, DMSO, r.t., 1 h, 26 %; (c) NaOH, MeOH, $H_2O$, 60 °C, 12 h, 49 %; (d) $RNH_2$, EDCI, DIEA, HOAt, DMF, -20 °C to r.t., overnight, 20%-52 %.

Although the synthetic route for cyclopropanation is shorter in **Scheme 1**, the yield is low (20-30 %). Thus, to prepare two selected fragments, we proposed another synthetic route modified from a reported sequence of reactions,[6] from which the yield reached 82 %. The synthesis of fragment **F47** is illustrated in **Scheme 2**. In order to remove protecting groups in the last step, strongly acidic condition was attempted firstly, but no desired compound was obtained. The high-resolution mass spectra (HRMS) of main product indicated that PMB groups were removed successfully, but the guanidine moiety unexpectedly formed a six-membered ring with triazole ring (Supplementary Figure 8). Alternatively, reductive hydrogeneration employing Pd/C catalyst was attempted to remove Cbz group, but no new product was detected. Then, we moved on to the synthesis of **F81**. When the protecting groups in compound **16** were attempted to remove under strongly acidic condition, HRMS of the main product again showed that

PMB groups were removed, whereas the guanidine moiety also formed a six-membered ring with pyrazole ring (Supplementary Figure 8).

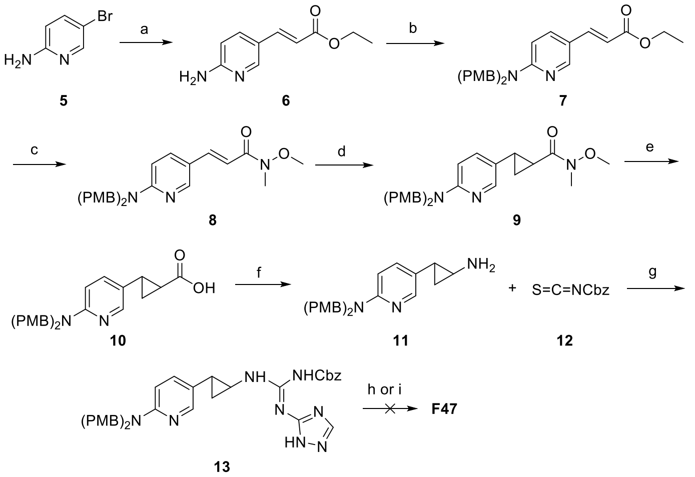


**Scheme 2.** Synthesis of F47. *Reagents and conditions*: (a) Ethyl acrylate, $Pd(OAc)_2$, DIEA, $P(o\text{-}tol)_3$, DMF, 100 °C, 20 h, 74 %; (b) PMB-Cl, NaH, DMF, 0 °C, 1 h, 71 %; (c) (i) 2M NaOH in $EtOH/H_2O$, r.t., 24 h; (ii) N,O-dimethylhydroxylamine hydrochloride, EDCI, DMAP, DCM, r.t., 2 h, 40 % over two steps; (d) $Me_3SOI$, NaH, DMSO, 0 °C to r.t., 4 h, 82 %; (e) KOH, $EtOH/H_2O$, r.t., 24 h, 62 %; (f) (i) DPPA, TEA, benzene, 80 °C, 1 h; (ii) $H_2O$, 80 °C, 30 min, 43 % over two steps; (g) (i) DCM, 0 °C to r.t., 4 h; (ii) 1,2,4-triazol-5-amine, EDCI, DIEA, DCM, 0 °C, 1h, then r.t.,10 h, 32 % over two steps; (h) TFA, DCM, r.t., 5 h; (i) 10 % Pd/C, MeOH, r.t., 24 h.

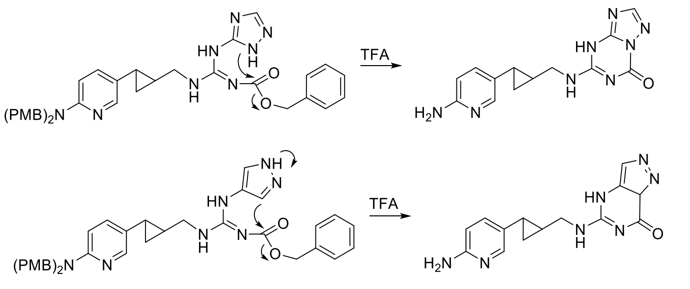


**Supplementary Figure 8.** Proposed formation of by-product indicated by HRMS.

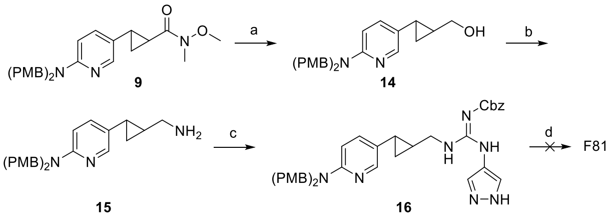


**Scheme 3.** Synthesis of F81. *Reagents and conditions*: (a) $LiAlH_4$, THF, 0 °C, 2 h, 72 %; (b) (i) MsCl, TEA, DCM, 0 °C, 2 h; (ii) $NaN_3$, DMF, 60 °C, 6 h; (iii) $PPh_3$, THF, $H_2O$, r.t., 12 h, 25 % over three steps; (c) (i) DCM, 0 °C to r.t., 4 h; (ii) pyrazol-4-amine, EDCI, DIEA, DCM, 0 °C, 1 h, then r.t.,10 h, 63 % over two steps; (d) TFA, DCM, r.t., 5 h.

### NMR spectra and HRMS of fragments F01-05

Spectra of **F01**

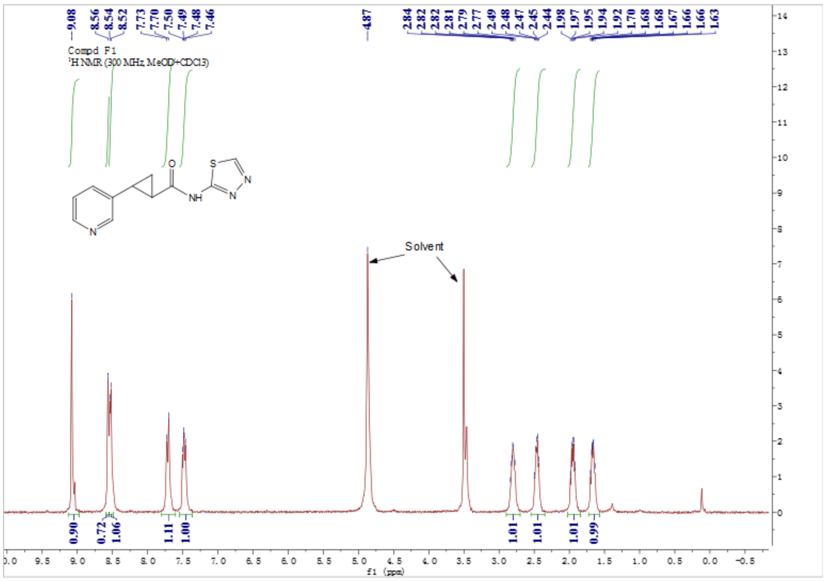


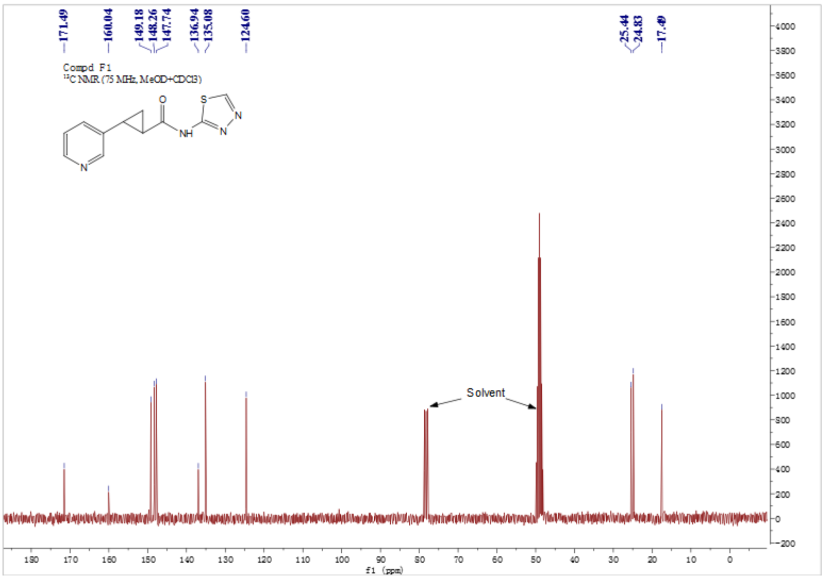

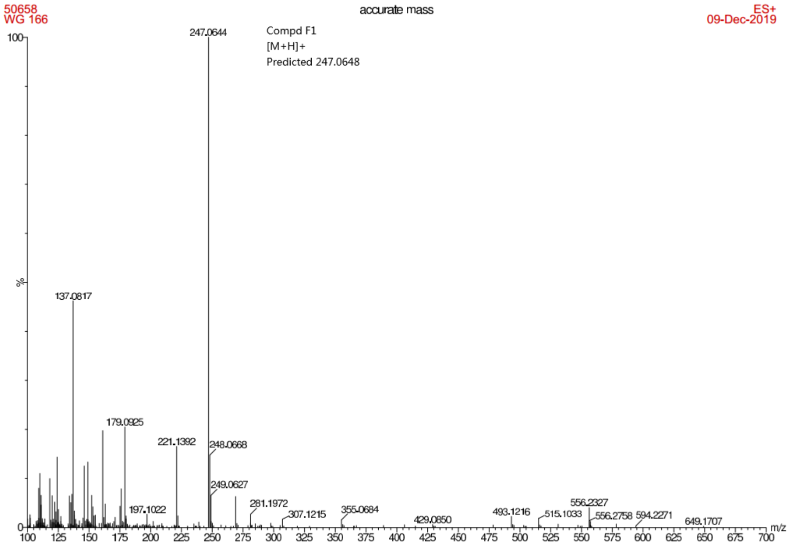


Spectra of **F02**

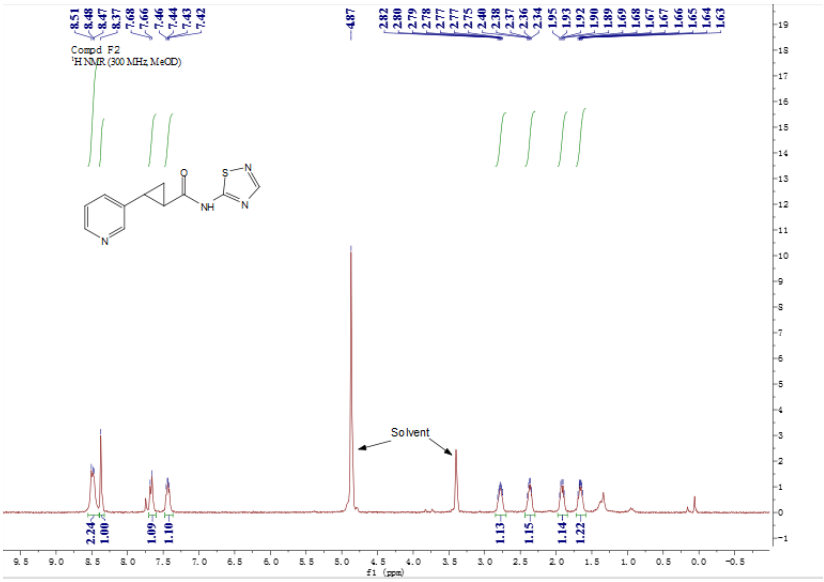

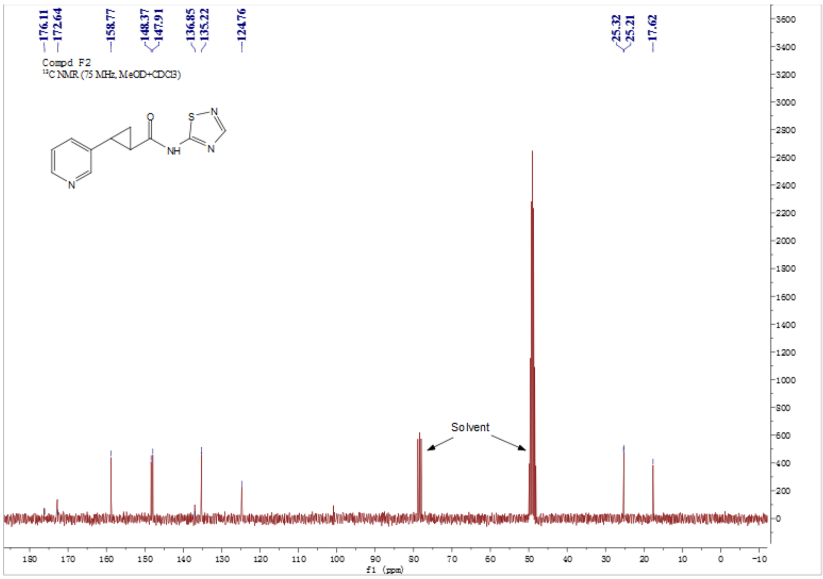

176.11
172.64
158.77
148.37
147.91
136.85
135.22
124.76
25.32
25.21
17.62
Compd F2
$^{13}$C NMR (75 MHz, MeOD+CDCl3)
Solvent
f1 (ppm)


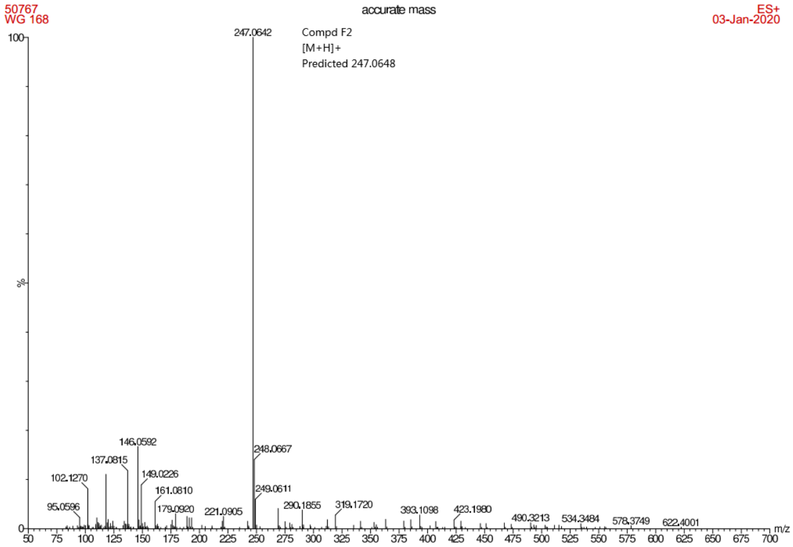

50767
WG 168
accurate mass
ES+
03-Jan-2020
247.0642
Compd F2
[M+H]+
Predicted 247.0648
146.0592
137.0815
102.1270
149.0226
161.0810
95.0596
179.0920
221.0905
248.0667
249.0611
290.1855
319.1720
393.1098
423.1980
490.3213
534.3484
578.3749
622.4001
m/z
%

Spectra of **F03**

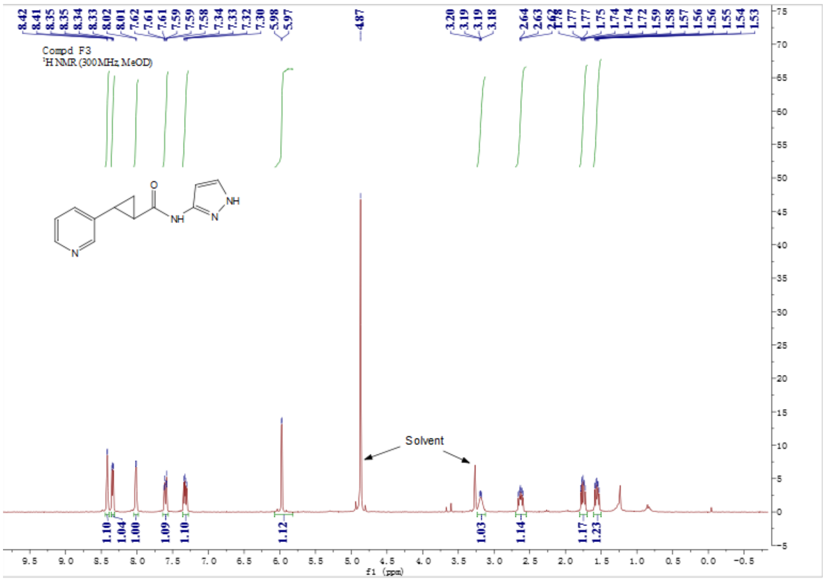


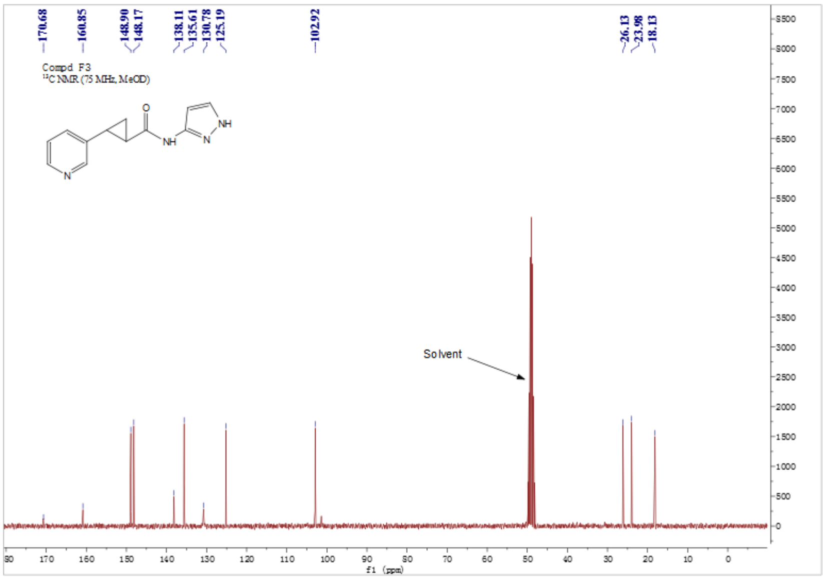

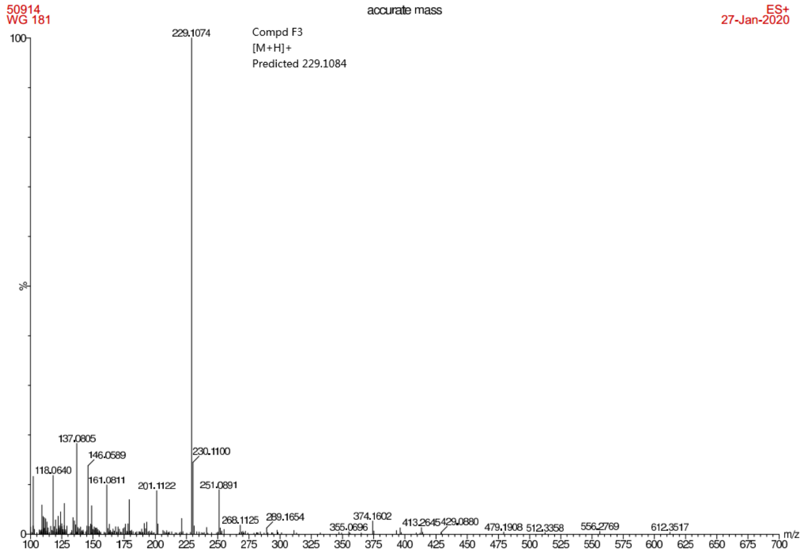


### Spectra of **F04**

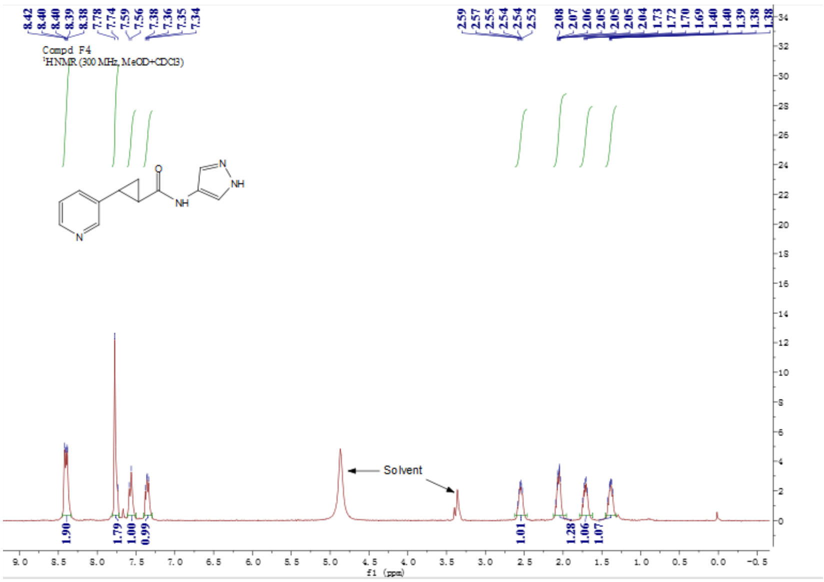

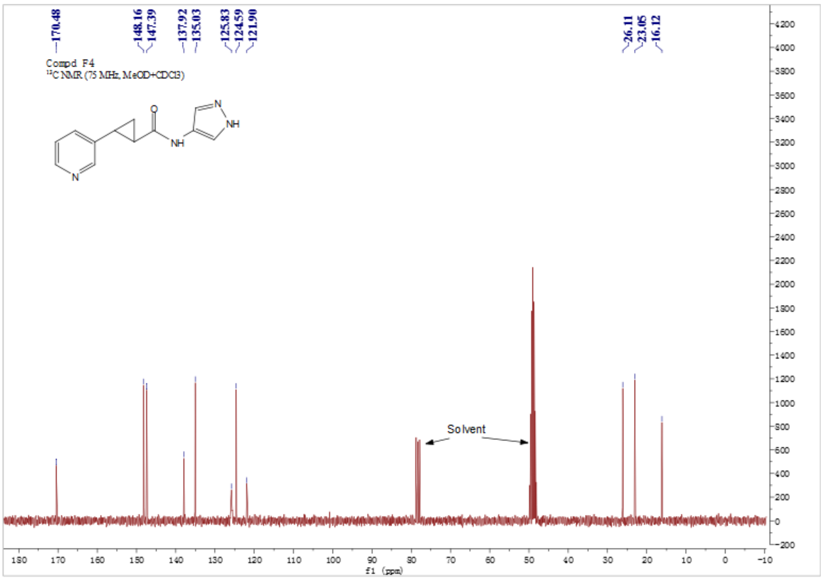
170.48
148.16
147.39
137.92
135.03
125.83
124.59
121.90
26.11
23.05
16.12
Compd F4
13C NMR (75 MHz, MeOD+CDCl3)
Solvent
f1 (ppm)

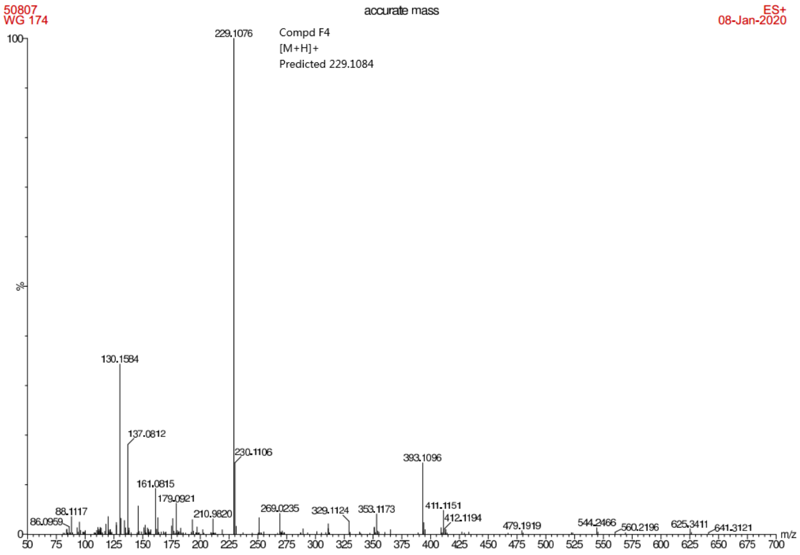
50807
WG 174
accurate mass
ES+
08-Jan-2020
229.1076
Compd F4
[M+H]+
Predicted 229.1084
130.1584
137.0812
230.1106
393.1096
161.0815
179.0921
88.1117
86.0959
210.9820
269.0235
329.1124
353.1173
411.1151
412.1194
479.1919
544.2466
560.2196
625.3411
641.3121
m/z
%

Spectra of **F05**

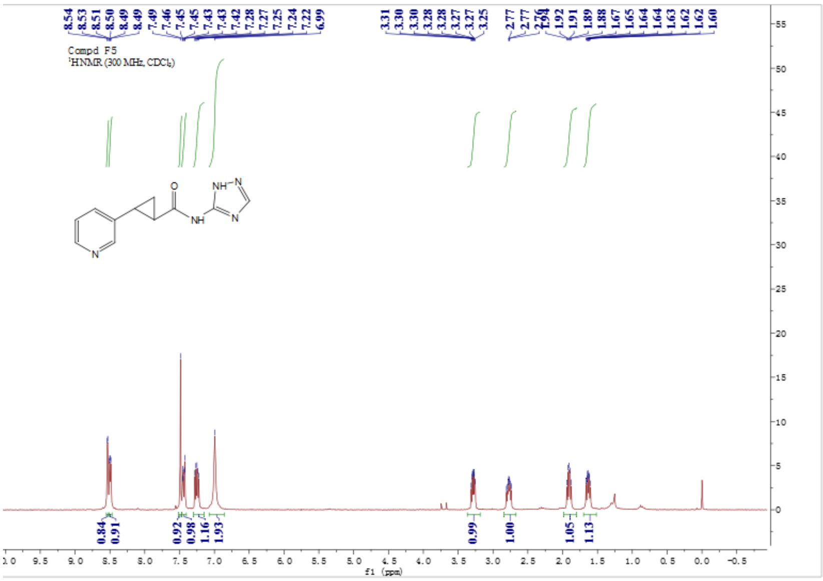


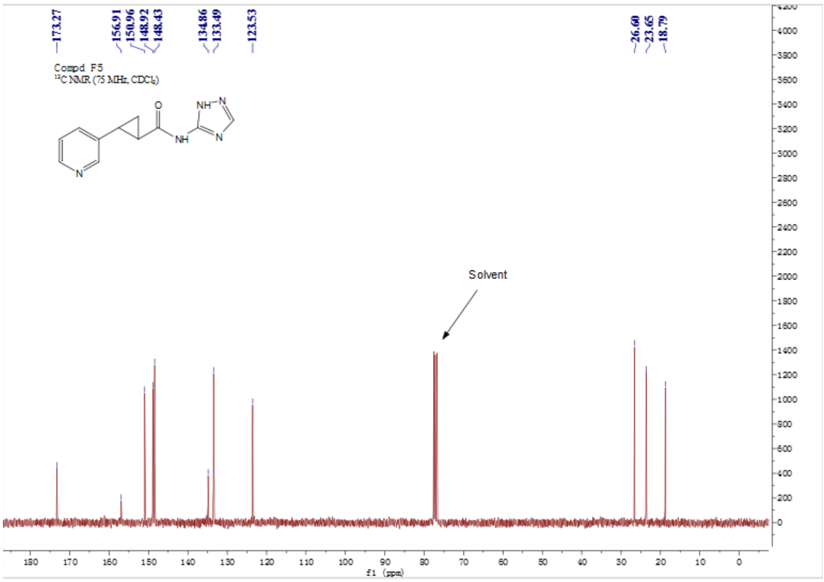

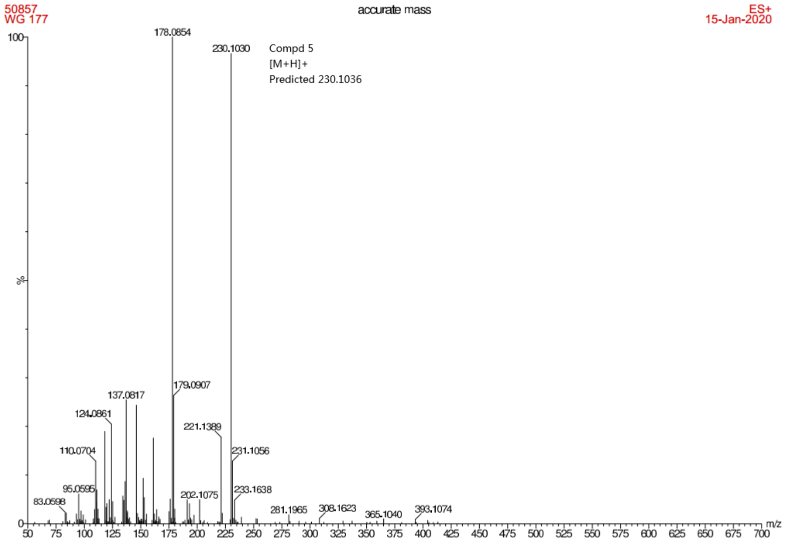

50857
WG 177
accurate mass
ES+
15-Jan-2020
Compd 5
[M+H]+
Predicted 230.1036
178.0854
230.1030
179.0907
137.0817
124.0861
221.1389
110.0704
231.1056
95.0595
233.1638
83.0598
202.1075
281.1965
308.1623
365.1040
393.1074
%
m/z
